# A shift in the pathotype diversity and complexity of *Phytophthora sojae*, causal agent of root and stem rot of soybean, in Brazil

**DOI:** 10.1101/2022.08.19.504165

**Authors:** I.C.A. Batista, M.P.C. Silva, A. L. Silva Junior, M.P. Gonzalez, M. P. Camargo, A. Figueiredo, B.T. Hora Junior, E.S.G. Mizubuti

## Abstract

Soybean root rot disease caused by the oomycete *Phytophthora sojae* is particularly destructive, once it is a host-specific pathogen that can infect and kill soybean plants at any stage of growth. The use of resistant cultivars is the most effective method of controlling the disease. Therefore, monitoring changes in the adaptability of *P. sojae* population to resistance genes (*Rps*) is extremely important not only from an epidemiological point of view but mainly for the management of the disease and durability of the *Rps* genes. We investigated the pathotypes of 40 isolates of *P. sojae*, sampled from the region with a higher incidence of soybean root rot in Brazil, using a set of 14 soybean differentials. The first study investigating pathotype diversity in Brazil was published in 2013. A decade later, we are reporting a major shift in pathotype diversity and complexity. This information can be useful for breeding programs aiming at developing soybean cultivars with resistance to root and stem rot and optimize the usage of genes and germplasm.

## Disease Note

Phytophthora root rot (PRR) of soybean, caused by the oomycete *Phytophthora sojae*, can significantly reduce yield and the magnitude of the economic losses ranked PRR among the top five diseases of this crop (Matthiesen et al. 2021). During the 2020/21 season, soybean seedling damping-off and root rot were observed in different municipalities in Rio Grande do Sul, Santa Catarina and Paraná states, Brazil. Soil samples from areas where symptomatic plants were reported were collected and analyzed using a leaf baiting technique and bioassays methods. Soil samples were placed into a 1000 mL-plastic container and flooded with 500 mL of distilled water for 24 h. Trifoliate leaves of the *P. sojae*-susceptible cultivar Williams were placed afloat on water and incubated at 23 ° C for up to 7 days. For the bioassay method, 20 to 30 seeds of susceptible soybean cv. Williams were distributed on the surface of infested soil samples and covered with a wet layer of planting substrate. After seed germination, the pots were flooded at every 48 h until seedlings with damping-off and water-soaked lesions on the hypocotyl were observed. Water-soaked lesions formed on the leaves or in the hypocotyl were excised using a scalpel, plated on *Phytophthora* selective medium (PBNIC) (Dorrance et al. 2008) and incubated at 25° C and 12 h-photoperiod. Cultures were characterized as *P. sojae* by observing morphological traits and by conventional PCR amplification using the *P. sojae*-specific primers PSOJF1 and PSOJR1 (Bienapfl et al. 2011). Forty isolates of *P. sojae* were obtained from 15 soybean fields distributed in 11 municipalities. Pure cultures were transferred to a slant of carrot-agar for storage at 18° C. Pathotypes of *P. sojae* isolates were determined using the hypocotyl technique on a set of 14 differential soybean genotypes: L88-8470 (*Rps*1a), L77-1863 (*Rps*1b), L75-3735 (*Rps*1c), L93-3312 (*Rps*1d), L77-1794 (*Rps*1k), L76-1988 (*Rps*2), L83-570 (*Rps*3a), L89-1541 (*Rps*3b), L92-7857 (*Rps*3c), L85-2352 (*Rps*4), L85-3059 (*Rps*5), L89-1581 (*Rps*6), L93-3258 (*Rps*7), PI 399073 (*Rps*8), and cultivar Williams (rps) as susceptible. Ten seedlings of each differential were inoculated by injecting approximately 0.1 mL of mycelial slurry of each isolate in the hypocotyl. The isolate Ps36.1 *P. sojae* (virulence formula 1b, 1d, 3a, 3b, 3c, 4, 5, 6, 7 and 8, pathotype 21773) was used as a positive control in this assay. The plants were incubated in a dew chamber at 24 ° C for 24 h and then placed in a greenhouse at 24°C for 5 days when they were assessed. The number of dead seedlings were recorded to score the reaction as susceptible (≥ 60% of dead seedlings) or resistant (< 60% of dead seedlings). The distribution of susceptible reactions, pathotype complexity, pathotype frequency, and diversity indices were estimated and the octal nomenclature was used to summarize pathotype descriptions as originally described (Dorrance et al. 2003). A total of 27 pathotypes out of 40 *P. sojae* isolates were identified. The most frequent pathotypes were 25573 representing 10% of the isolates, followed by the pathotypes 27573, 27771, 27773 and 77573 (all corresponding to 7.50% of the isolates) and pathotypes 27571 and 27761 (5% of the isolates). The other 50% (20 isolates) were single pathotypes. All *P. sojae* isolates were virulent on *Rps*1d (Supplementary Table 1, Supplementary Figure 1). Over 60% of the isolates were virulent on *Rps*1b, *Rps*1k, *Rps*2, *Rps*3a, *Rps*3c, *Rps*4, *Rps*5, *Rps*6 and *Rps*7. Twenty isolates were virulent to *Rps*8 (50%), 14 isolates (35%) were virulent to *Rps*3b, eight isolates (20%) were virulent on *Rps*1a, while only seven isolates (17.50%) were virulent on *Rps*1c. All *Rps* genes were overcome by at least one isolate (Supplementary Figure 2). The mean number of resistance genes that a given pathotype could overcome (complexity) was 9.65. The calculated diversity indices were: Simple diversity = 0.67; Gleason = 7.05; Shannon = 3.15; Simpson diversity = 0.95; and Evenness = 0.96. There is a tendency towards greater variability of *P. sojae* in the current population when compared with the results of a previous study (Costamilan et al., 2012). Furthermore, there has been an important shift towards greater complexity of pathotypes over time. The average complexity reported by Costamilan et al. (2012) with 37 isolates from the same region (South) was 6.70 while for the 2020/2021 season the estimated complexity was 9.65. Compared to recent results from studies conducted in The United States, the Brazilian population of *P. sojae* seems to be more variable and the changes in the avirulence gene pool seem to occur at a faster pace. This scenario imposes the need for special attention of the breeding programs aimed at controlling soybean PRR in Brazil, as well as for the strategic deployment of the resistance genes currently known and largely used by several seed companies.

## Acknowledgements

We acknowledge Bayer Crop Science for financial support (Convênio Bayer-UFV No 271/2019).

The authors declare that they have no conflict of interest.

## Notes

### Competing Interest Statement

The authors have declared no competing interest.

