## Supplemental table 1 and Supplemental figures 1 and 2 for "A shift in the pathotype diversity and complexity of *Phytophthora sojae*, causal agent of root and stem rot of soybean, in Brazil"

### Supplementary Material

**Supplementary Table 1.** Virulence formulae and octal code of the pathotypes of *Phytophthora sojae* isolated from soil samples in the South Region in Brazil.

| Pathotype | Number of isolates | Municipality | Octal |
|---|---|---|---|
| 1b, 1d, 2, 3a, 3c, 4, 5, 6, 7, 8 | 4 | Tapes (2 isolates), Senador Salgado Filho (2) | 25573 |
| 1a, 1b, 1c, 1d, 1k, 2, 3a, 3c, 4, 5, 6, 7, 8 | 3 | Caçapava do Sul (2), Cruz Alta (1) | 77573 |
| 1b, 1d, 1k, 2, 3a, 3b, 3c, 4, 5, 6, 7 | 3 | Santo Augusto (2), Sarandi (1) | 27771 |
| 1b, 1d, 1k, 2, 3a, 3b, 3c, 4, 5, 6, 7, 8 | 3 | Não-Me-Toque (1), Sarandi (2) | 27773 |
| 1b, 1d, 1k, 2, 3a, 3c, 4, 5, 6, 7, 8 | 3 | Não-Me-Toque | 27573 |
| 1b, 1d, 1k, 2, 3a, 3b, 3c, 5, 6, 7 | 2 | Santo Augusto | 27761 |
| 1b, 1d, 1k, 2, 3a, 3c, 4, 5, 6, 7 | 2 | Abelardo Luz (1), Não-Me-Toque (1) | 27571 |
| 1a, 1b, 1c, 1d, 1k, 2, 3a, 3b, 3c, 4, 5, 6, 7 | 1 | Guarapuava | 77771 |
| 1a, 1b, 1c, 1d, 1k, 2, 3a, 3b, 3c, 4, 5, 6, 7, 8 | 1 | Guarapuava | 77773 |
| 1a, 1b, 1c, 1d, 1k, 2, 3a, 6 | 1 | Guarapuava | 77140 |
| 1a, 1b, 1c, 1d, 1k, 2, 5, 6, 7, 8 | 1 | Cruz Alta | 77363 |
| 1a, 1d, 3a, 5, 6 | 1 | Cruz Alta | 11160 |
| 1b, 1d, 1k, 2, 3a, 3b, 3c, 6, 7 | 1 | Sarandi | 27741 |
| 1b, 1d, 1k, 2, 3a, 3b, 6, 7 | 1 | Santo Augusto | 27341 |
| 1b, 1d, 1k, 2, 3a, 3c, 5, 6, 7 | 1 | Abelardo Luz | 27561 |
| 1b, 1d, 1k, 2, 3a, 4, 5, 6, 7 | 1 | Sarandi | 27171 |
| 1b, 1d, 2, 3a, 3b, 3c, 4, 5, 6, 7, 8 | 1 | Dom Pedrito | 25773 |
| 1b, 1d, 2, 3a, 3b, 4, 5, 6 | 1 | Dom Pedrito | 25370 |
| 1b, 1d, 2, 3a, 3c, 4, 5, 6 | 1 | Tapes | 25573 |
| 1b, 1d, 2, 3a, 3c, 4, 5, 7, 8 | 1 | Alegrete | 25533 |
| 1b, 1d, 2, 3a, 3c, 4, 8 | 1 | Senador Salgado Filho | 25512 |
| 1b, 1d, 2, 3a, 4, 6 | 1 | Senador Salgado Filho | 25150 |
| 1d, 1k, 2, 3a, 4 | 1 | Sarandi | 27171 |
| 1d, 2, 3a, 3c, 4, 5, 6, 7, 8 | 1 | Tapes | 25573 |
| 1d, 2, 3a, 3c, 4, 7 | 1 | Cruz Alta | 05511 |
| 1d, 2, 3c, 4, 5, 6, 7, 8 | 1 | Cruz Alta | 05473 |
| 1d, 2, 7 | 1 | Cruz Alta | 05001 |

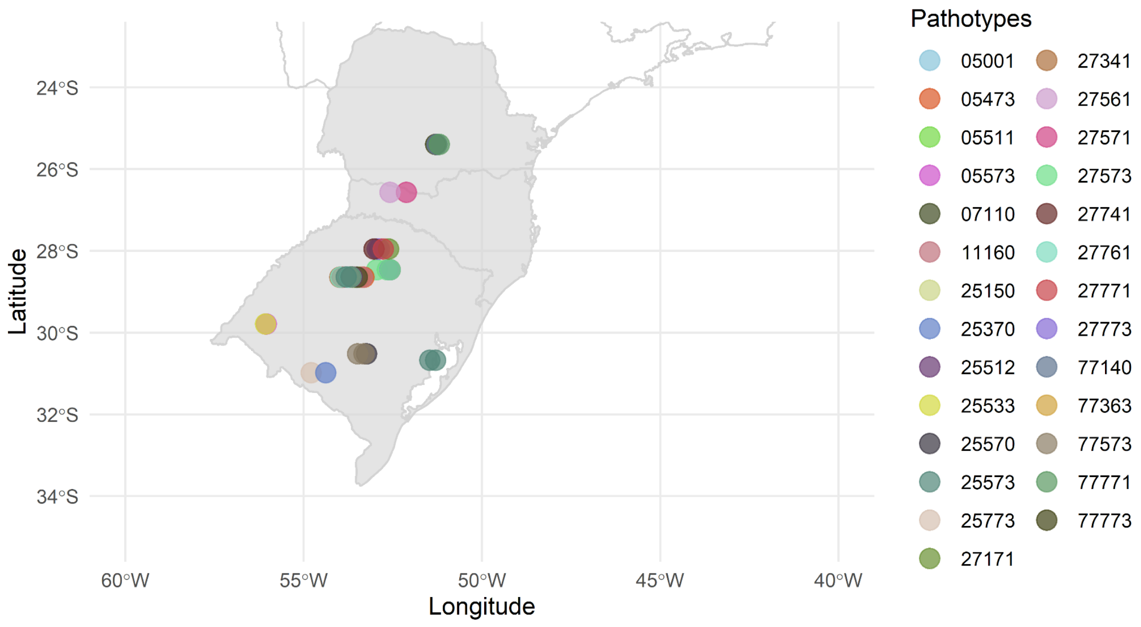


**Supplementary Figure 1**. Distribution of pathotypes of *Phytophthora sojae* recovered in a survey of soybean fields in Rio Grande do Sul, Santa Catarina and Paraná states, Brazil in the 2020/2021 crop season.

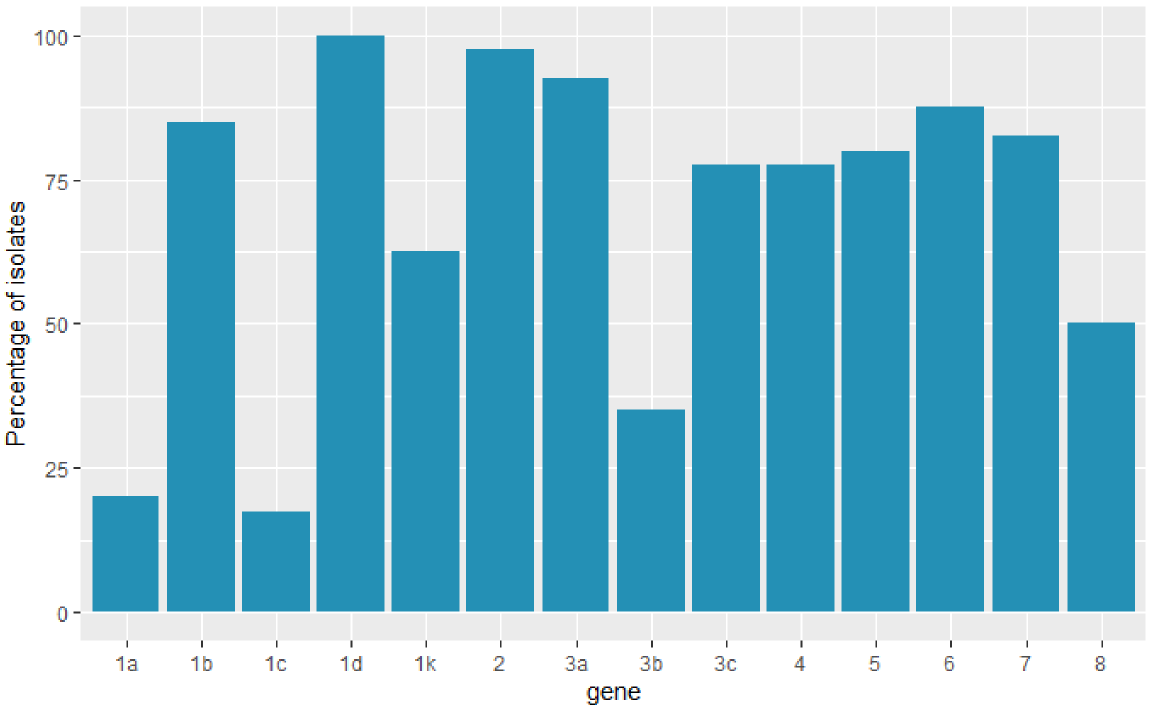


**Supplementary Figure 2**. Percentage of *Phytophthora sojae* isolates recovered in a survey of soybean fields in the South Region of Brazil in the 2020/2021 crop season, that were virulent on a specific *Rps* gene.
